# Proximity proteomics provides a new resource for exploring the function of Afadin and the complexity of cell-cell adherens junctions

**DOI:** 10.1101/2024.11.07.622507

**Authors:** Wangsun Choi, Dennis Goldfarb, Feng Yan, M. Ben Major, Alan S. Fanning, Mark Peifer

## Abstract

The network of proteins at the interface between cell-cell adherens junctions and the actomyosin cytoskeleton provides robust yet dynamic connections that facilitate cell shape change and motility. While this was initially thought to be a simple linear connection via classic cadherins and their associated catenins, we now have come to appreciate that many more proteins are involved, providing robustness and mechanosensitivity. Defining the full network of proteins in this network remains a key objective in our field. Proximity proteomics provides a means to define these networks. Mammalian Afadin and its Drosophila homolog Canoe are key parts of this protein network, facilitating diverse cell shape changes during gastrulation and other events of embryonic morphogenesis. Here we report results of several proximity proteomics screens, defining proteins in the neighborhood of both the N- and C-termini of mammalian Afadin in the premier epithelial model, MDCK cells. We compare our results with previous screens done in other cell types, and with proximity proteomics efforts with other junctional proteins. These reveal the value of multiple screens in defining the full network of neighbors and offer interesting insights into the overlap in protein composition between different epithelial cell junctions.

**Summary Statement:** Afadin BioID reveals new adherens junction proteins.

## Introduction

The multiprotein complexes at cell-cell and cell-extracellular matrix junctions establish and maintain epithelial tissue architecture and facilitate cell shape changes and cell migration during both morphogenesis and tissue homeostasis (Perez-Vale and Peifer, 2020). To accomplish these tasks, junctional protein complexes need to be connected to the contractile actomyosin cytoskeleton (Cronin and DeMali, 2021). Defining the full set of molecular components at each junction and unraveling their collective functions is a key challenge for our field.

Work over several decades revealed that cell-extracellular matrix junctions link to the cytoskeleton via a complex, layered network of dozens of proteins (Case and Waterman, 2015). In contrast, until the mid-2000’s, the view of cytoskeletal connections at cell-cell adherens junctions (AJs) was much simpler, suggesting direct linkage (Gates and Peifer, 2005). Transmembrane classic cadherins linked cells to one another by homophilic interactions. Beta-catenin bound both the cadherin cytoplasmic tail and alpha-catenin, and alpha-catenin then bound actin. However, this simplistic picture of junction:cytoskeletal linkage has been replaced by one that is more complex in several ways. First, it is now clear that a large network of proteins is localized to AJs. Many are large, multidomain proteins that interact with one another by complex, multivalent linkages (Rouaud et al., 2020). Second, AJs are mechanosensitive and mechanoresponsive (Buckley et al., 2014; Campas et al., 2024; Wang et al., 2022), with conformational change and protein recruitment strengthening cytoskeletal connections when junctions are under mechanical tension (Yap et al., 2018).

Many additional proteins are now known to be part of this network (Perez-Vale and Peifer, 2020), including p120, Par3, Afadin/Canoe, ZO-1 family members, vinculin, and Ajuba. Specialized proteins are enriched at tricellular junctions where three cells meet, including vertebrate tricellulin, angulin-1/ LSR, angulin-2/ILDR1 and angulin-3/ ILDR2, and Drosophila Anakonda, Gliotactin, Sidekick and M6 protein (Higashi and Chiba, 2020). These need to link to the core cadherin-catenin complex (van den Goor and Miller, 2022). In addition to actin and non-muscle myosin, additional cytoskeletal regulators are also enriched at AJs, including Eva/VASP proteins (Gates et al., 2007), and at least in some cell types, Diaphanous-class formins (Homem and Peifer, 2008) and the Arp2/3 complex (Kovacs et al., 2002). Signaling proteins also localize to AJs, including both transmembrane receptors like the Epidermal Growth Factor Receptor (Fedor-Chaiken et al., 2003) and cytoplasmic signaling molecules, like non-receptor tyrosine kinases in the Src (Calautti et al., 1998) and Abl families (Stevens et al., 2008).

We focus on the roles of Afadin and its Drosophila homolog Canoe in junction-cytoskeletal linkage. They play important roles in cell shape change and cell rearrangements during events ranging from initial positioning of cell-cell AJs (Choi et al., 2013) to the apical constriction and convergent elongation events of gastrulation (Ikeda et al., 1999; Sawyer et al., 2011; Sawyer et al., 2009; Zhadanov et al., 1999) to later collective cell migration (Boettner et al., 2003). Both also are important for the architecture of adult tissues, ranging from fly eyes (Gaengel and Mlodzik, 2003; Miyamoto et al., 1995) to mammalian kidneys (Yang et al., 2013). Unlike the cadherin-catenin complex, which localizes all along the lateral cell membrane, Afadin/Canoe is more tightly enriched at the apical end of the lateral interface, in the structure often referred to as the zonula adherens, though in at least one mammalian cell type, Afadin moves to an even more apical level as AJs mature. In this tissue it moves apically from a location mixed with cadherin-catenin complexes, but still remains basal to the tight junctions (Mangeol, 2024).

Afadin/Canoe family members are found across the animal kingdom. They share a complex, multidomain protein architecture (Fig. 1A; Gurley et al., 2023; Smith, 2023). Most N-terminal are two RA domains, known to bind small GTPases in the Ras/Rap1 family. These GTPases use Afadin/Canoe as effectors, activating them upon binding. Next follow three additional conserved protein domains: a Forkhead-associated (FHA) domain, known in other proteins to binding phosphorylated peptides, a Dilute domain, only known from Afadin/Canoe and the non-conventional Myosin V family, and a PSD-95/discs large/zona occludens (PDZ) domain. This set of conserved protein domains is followed by a long intrinsically disordered region that includes a region or regions that bind filamentous actin. Afadin/Canoe’s multidomain architecture allows it to interact with multiple proteins. These include small GTPases in the Ras/Rap1 family that bind the RA domains (McParland et al., 2024; Smith, 2023), the ADIP protein that can bind the Dilute domain (Asada et al., 2003), Scribble, which can bind the FHA domain (Goudreault et al., 2022), PLEKHA7, which can bind both RA and PDZ domains (Kurita et al., 2013), and the transmembrane junctional proteins Nectins (Takahashi et al., 1999), E-cadherin (Sawyer et al., 2009), Neurexin (Zhou et al., 2005), and JAM-A (Ebnet et al., 2000), which along with Eph family receptors (Buchert et al., 1999), the Notch ligand Jagged (Popovic et al., 2011), and the kinase/GEF/GAP Bcr (Radziwill et al., 2003) can bind the PDZ domain. Actin (Mandai et al., 1997), alpha-catenin (Pokutta et al., 2002), ZO-1 (Takahashi et al., 1998), Ponsin (Mandai et al., 1999), Lgn (Carminati et al., 2016), and profilin (Boettner et al., 2000) can bind to sites in the intrinsically disordered region. A recent cryo-EM structure revealed cooperative interactions between a conserved motif in the intrinsically disordered region that binds both actin and the actin-binding domain of alpha-catenin (Gong et al., 2024). Given this diversity of interaction domains and regions, it seems likely additional Afadin binding partners exist.

**Fig. 1.**
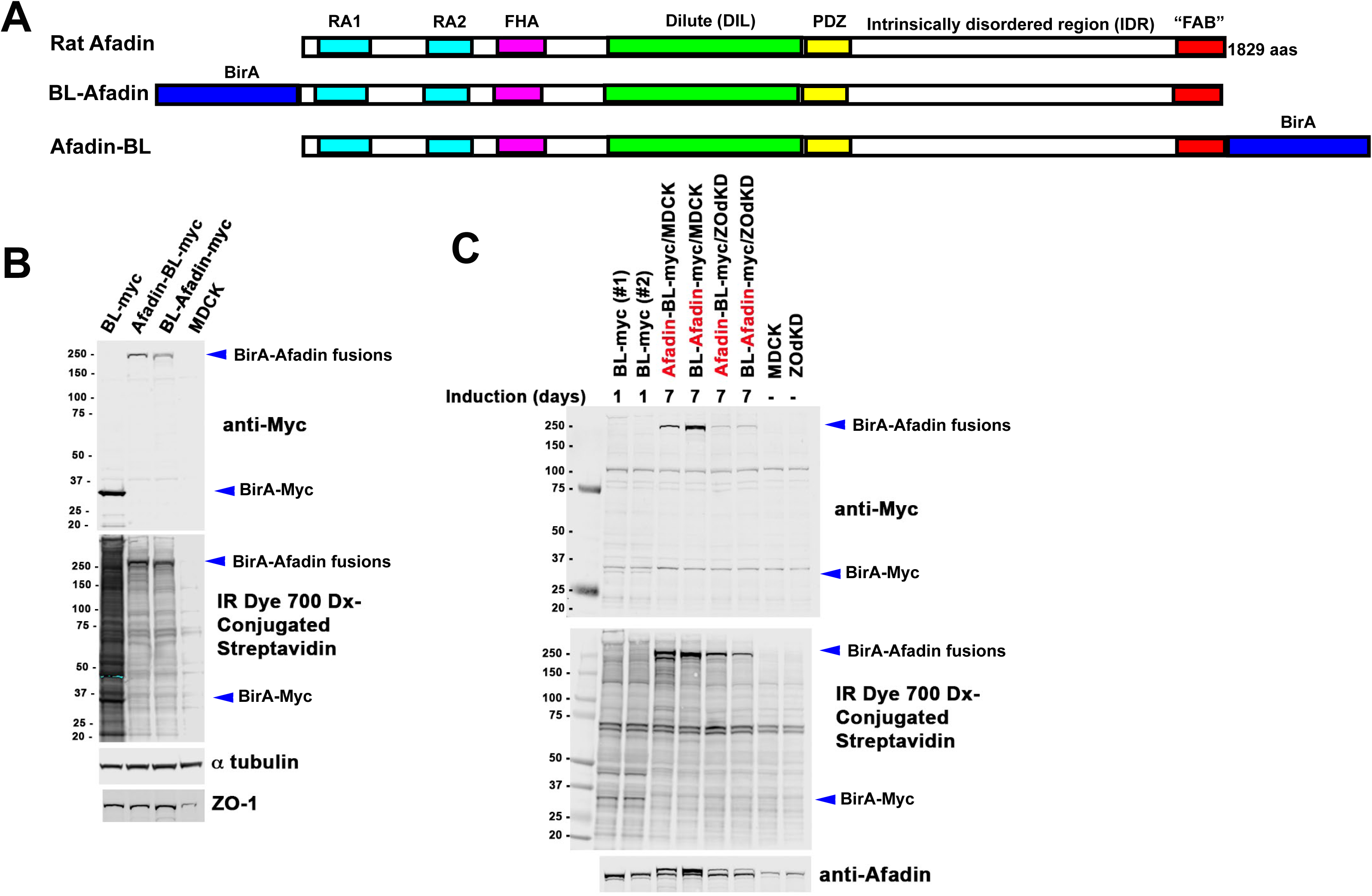
Designing and validating reagents to identify potential Afadin protein partners and neighbors. (A) Diagram of Afadin and the BirA*-fusions used. (B) Immunoblot of cell extracts from MDCK cells induced to express our BirA*-myc control, Afadin-BL-myc, BL-Afadin-myc or no construct. Top panel. Immunoblot with anti-myc antibody to detect expressed constructs. Middle-panel. Blot with IR Dye 700 Dx-Conjugated Streptavidin to detect biotinylated proteins. Tubulin serves as a loading control. (C) Immunoblot of cell extracts of MDCK cells or ZO-knockdown MDCK cells induced to express our BirA*-myc control, Afadin-BL-myc, BL-Afadin-myc or no construct. Induction time of the BirA*-myc control was only one day, while induction time for the others was seven days. Top panel. Immunoblot with anti-myc antibody to detect expressed constructs. Middle-panel. Blot with IR Dye 700 Dx-Conjugated Streptavidin to detect biotinylated proteins. Bottom panel. Immunoblot with anti-Afadin antibodies to confirm expression of tagged proteins.

One key knowledge gap in our field is the full identity of the network of proteins connecting AJs to the cytoskeleton. Proteomics tools can help fill this gap. Proximity-labeling approaches like APEX or BioID offer the opportunity to take an assumption-free approach to identifying proteins in this network (Bosch et al., 2021). By fusing the protein of interest to an engineered enzyme that can add biotin to nearby proteins, and then purifying biotinylated proteins using streptavidin, one can identify both direct and indirect interactors (Roux et al., 2018). We used this approach to define and contrast proteins in the neighborhood of either the N-or C-termini of mammalian Afadin in one of the best-characterized of all mammalian epithelial cell lines, MDCK cells.

## Results

### Designing and validating reagents to identify potential Afadin protein partners and neighbors

Afadin and its Drosophila homolog Canoe are conserved multidomain proteins that are key components in the network of proteins linking cadherin-based AJs to the actomyosin cytoskeleton. Some domains or regions have known binding partners, but the interactors for other well conserved domains remain unknown. We sought an unbiased approach to identify proteins within the Afadin protein network, including those binding directly and those that are proximal. To do so, we used BioID-based proximity labeling, in which the promiscuous biotin ligase enzyme (BirA-R118G, referred to as BirA*) is fused a protein of interest. Following the addition of exogenous biotin to live cells expressing a BirA*:bait fusion, proteins within a ∼10-50 nm sphere are biotinylated. Biotinylated proximal proteins are then purified with streptavidin for downstream mass spectroscopy identification and quantification (Sears et al., 2019)).

We previously cloned the coding sequence of mammalian Afadin tagged with both BirA* and a myc-epitope at either the N- or C-terminus (BL-Afadin or Afadin-BL) into doxycycline-regulated vectors (Fig. 1A; Bonello et al., 2019). In our previous analyses of Afadin function, we used the well-characterized epithelial cell line MDCK (Choi et al., 2016). We thus generated stable MDCK cell lines in which each fusion protein– BL-Afadin or Afadin-BL– was expressed after doxycycline-withdrawal (Bonello et al., 2019), alongside a control cell line expressing BirA* alone. We had also used MDCK cells in which both ZO-1 and ZO-2 were knocked down—in these cells junctional architecture is dramatically altered, with elevated levels of Afadin and the assembly of a highly contractile sarcomeric actomyosin cytoskeleton at AJs (Choi et al., 2016). We thus also created stable lines in for each fusion protein in this cell background.

We first assessed the expression of proteins, using antibodies to the myc-epitope. MDCK cells transfected with the BirA* construct alone, with BL-Afadin, or with Afadin-BL, were incubated in medium containing biotin. Immunoblotting revealed myc-tagged proteins of the expected sizes (Fig. 1B, top panel). In parallel, we transfected ZO-knockdown cells with the same constructs, and once again observed myc-tagged bands consistent with the sizes of BL-Afadin or Afadin-BL (Fig 1C, top panel). Re-probing this blot with antibodies to Afadin confirmed these are myc-tagged Afadin, as we observed doublets of wildtype and tagged Afadin in cells in which our constructs were expressed (Fig.1C, bottom panel). Finally, we re-probed these blots with IR Dye 700 Dx-Conjugated Streptavidin, which binds tightly to any biotinylated protein. We observed biotinylated proteins of the size of our constructs (Fig. 1B, C, middle panels), and also saw other biotinylated proteins. MDCK cells expressing BirA* alone had much higher levels of biotinylated proteins relative to the BL-Afadin or Afadin-BL cells, when cells were incubated for the same time (Fig. 1B, middle panel), and had similar levels of overall biotinylated protein when incubated with biotin for much shorter periods (1 versus 7 days; Fig. 1C, middle panel).

### BirA*-tagged versions of Afadin localize to cell-cell junctions

To assess whether tagging altered protein localization, we examined whether the myc-tagged BirA*/Afadin fusions localized to cell-cell junctions, like wildtype Afadin. To do so, we immunostained cells with antibodies to Afadin, the junctional protein ZO-1, F-actin, and, in the same cells, visualized the localization of biotinylated proteins by immunostaining with an Alexa-568 conjugated streptavidin. Consistent with specificity, the streptavidin signal was highly enriched at cell-cell junctions and co-localized with BL-Afadin (Fig. 2B-B’”) or Afadin-BL (Fig.2D-D’”). In contrast, in cells expressing BirA* alone, the streptavidin signal filled the whole cell, with no apparent enrichment at cell junctions (Fig. 2F-F’”). In each case, the streptavidin signal was dependent on addition of exogenous biotin (Fig 2A’ vs B’, C’ vs D’, E’ vs F’).

**Fig. 2.**
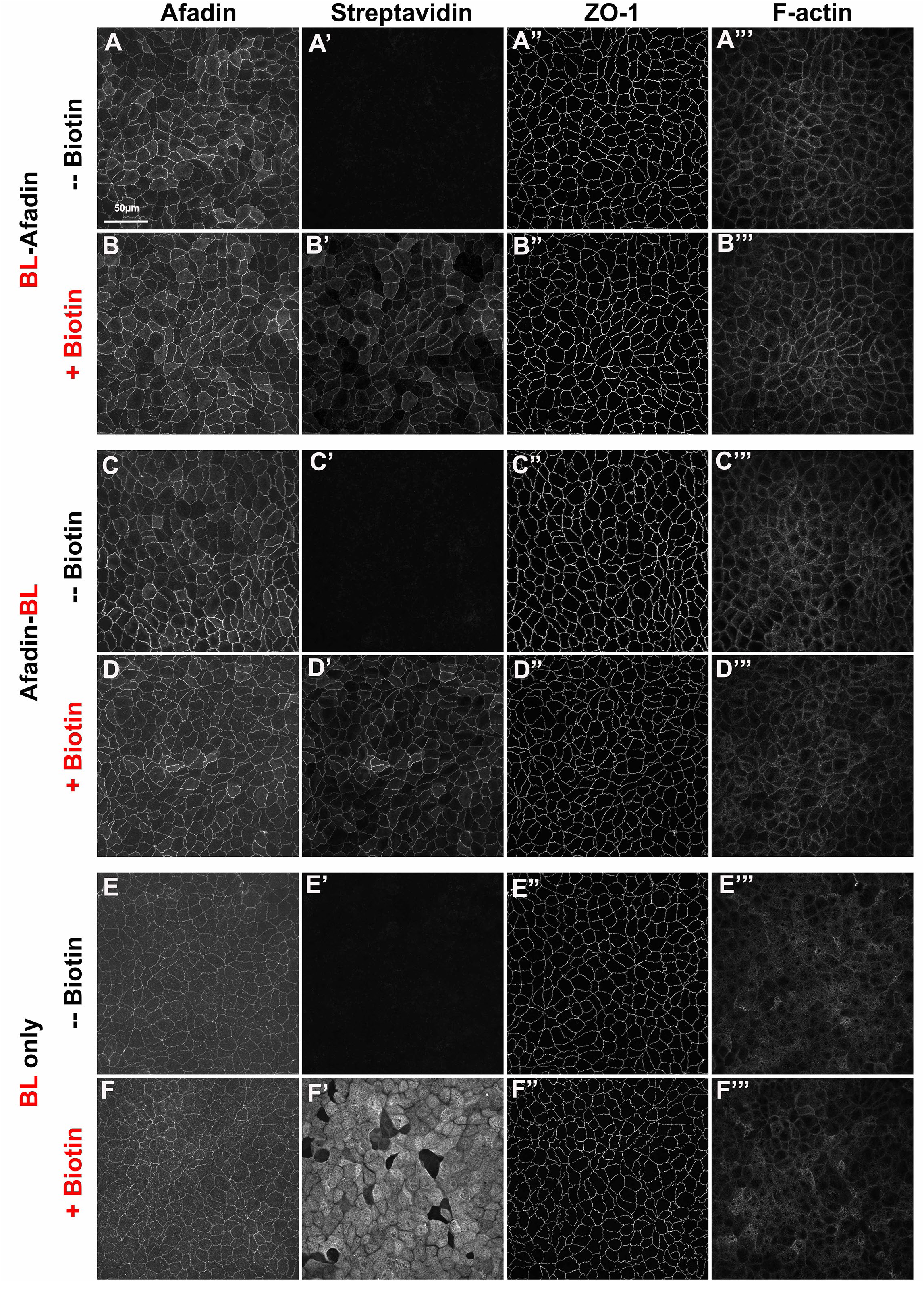
The BirA*-tagged versions of Afadin localize to cell junctions. MDCK cells cultured to confluence and prepared for immunofluorescence. Cells were cultured without or with added biotin in the medium. (A,B) Stable cell line expressing BL-Afadin. (C,D) Stable cell line expressing Afadin-BL. In both cases, cells with added biotin accumulate a streptavidin-labeled protein co-localizing with Afadin at cell junctions. (E,F) Stable cell line expressing our BirA*-myc control. In this case, addition of biotin leads to accumulation of streptavidin-labeled proteins throughout the cell.

### Some junctional proteins are biotinylated in cells expressing BirA*-tagged versions of Afadin

To further verify localization of our fusions to cell-cell junctions, and to begin to explore proteins that are proximal to Afadin, we examined selected junctional proteins. We incubated cells expressing either Afadin-BL or BL-Afadin in biotin over a time course of hours. We then affinity purified biotinylated proteins from cells using streptavidin before immunoblot analysis of selected junctional and cytoskeletal proteins (Fig. 3A). The tight junction proteins ZO-1 and ZO-2, known Afadin binding partners (Yamamoto et al., 1997), were readily recovered in the streptavidin pull down in cells expressing either Afadin-BirA* fusion (Fig. 3A, top two panels). We also recovered the core AJ proteins E-cadherin and beta-catenin (Fig. 3A, panels 3 and 4), though recovery of beta-catenin was weaker in cells expressing BL-Afadin. The heavy chain of cytoplasmic myosin was also detected (Fig. 3A, panel 5). However, not all junctional proteins were enriched—for example the tight junction protein JAM was not detected (Fig. 3A, bottom panel).

**Fig. 3.**
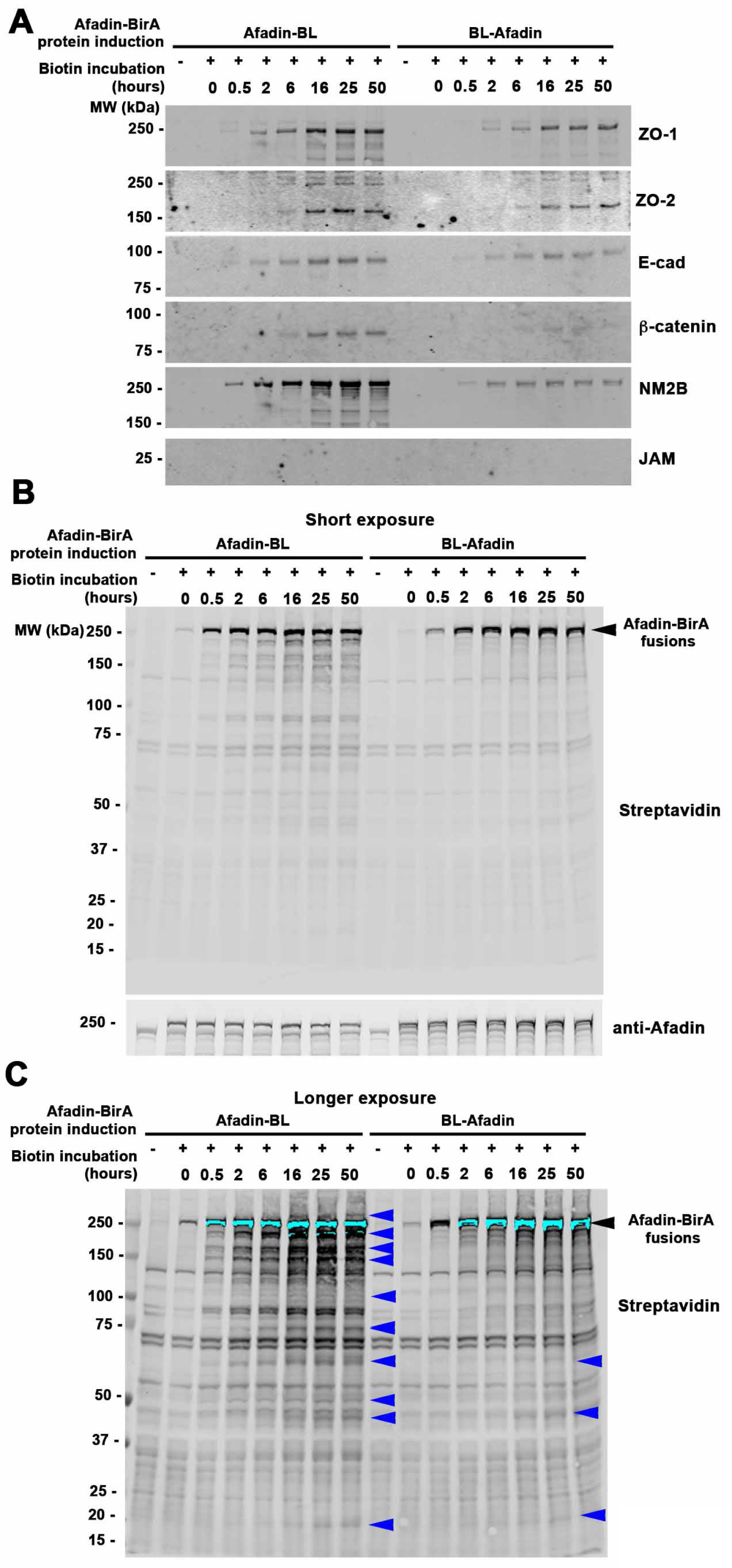
A subset of junctional proteins are biotinylated in cells expressing BirA*-tagged Afadin and a time course of accumulation of biotinylated proteins. (A) Stable MDCK cell lines carrying constructs expressing BirA*-tagged Afadin were cultured to confluence, exposed to biotin for varying amounts of time. Cell extracts were made, biotinylated proteins were affinity-purified from cells using streptavidin, analyzed by immunoblotting with antibodies to various junctional proteins. (B) Short exposure of an immunoblot of cell extracts from MDCK cells induced to express Afadin-BL-myc or BL-Afadin-myc, incubated with biotin over a time course of up to 50 hours, and blotted with IR Dye 700 Dx-Conjugated Streptavidin to detect biotinylated proteins, or with Afadin to detect the expressed fusion. The BirA*-tagged constructs are seen at all time points, with a low level of self-biotinylation without biotin addition and increasing levels beginning at 0.5 hours. (C) Long exposure. Other biotinylated proteins begin to be seen accumulating as the time course progresses (blue arrows).

### Mass spectrometry-based proximity proteomic analysis of Afadin identified 144 proximal proteins in MDCK cells

A biotin time course immunoblot analysis revealed robust biotinylation of our tagged Afadin constructs after short incubation (Fig. 3B), with biotinylation of Afadin-proximal proteins increasing at longer times of biotin treatment (Fig. 3B vs. C, blue arrows). Based on these data, we selected 24 hours of biotinylation for unbiased MS analysis. MDCK or ZOdKD cells expressing Afadin-BL, BL-Afadin, or the naked control BirA* fusion constructs were maintained in the presence of doxycycline to repress the expression of the transgenes (Tet-Off). For large-scale purification of biotinylated proteins, cells were cultured in 150 mm culture dishes in the presence of doxycycline until a monolayer formed (∼3-5 days). Once confluent, the culture was switched to doxycycline-free media to induce the fusion proteins and 50 μM biotin was added to the culture media for 24 hours. Cell lysates were then used for subsequent streptavidin purification, protein digestion with trypsin and label-free liquid chromatography–tandem MS analysis.

Biological duplicate experiments were performed for MDCK cells expressing the control BirA*, Afadin-BL, and BL-Afadin. A single replicate of Afadin-BL and BL-Afadin in ZOdKD cells was analyzed. We first used the SAINT (significance analysis of interactome) express algorithm to identify statistically significant discoveries versus control using a cutoff of SAINT ≥0.9. We then ordered the proteins on each list by the difference in average log2 LFQ intensity values between experimental and control (BirA* alone) groups. Doing so, we identified 144 Afadin proximal proteins. These included 95 proteins for BL-Afadin (Table 1; in each case excluding Afadin) and 106 proteins for Afadin-BL (Table 2). 55 proteins were shared on both lists (Table 3; Full proteomics data is in Table S1).

**Table 1.**
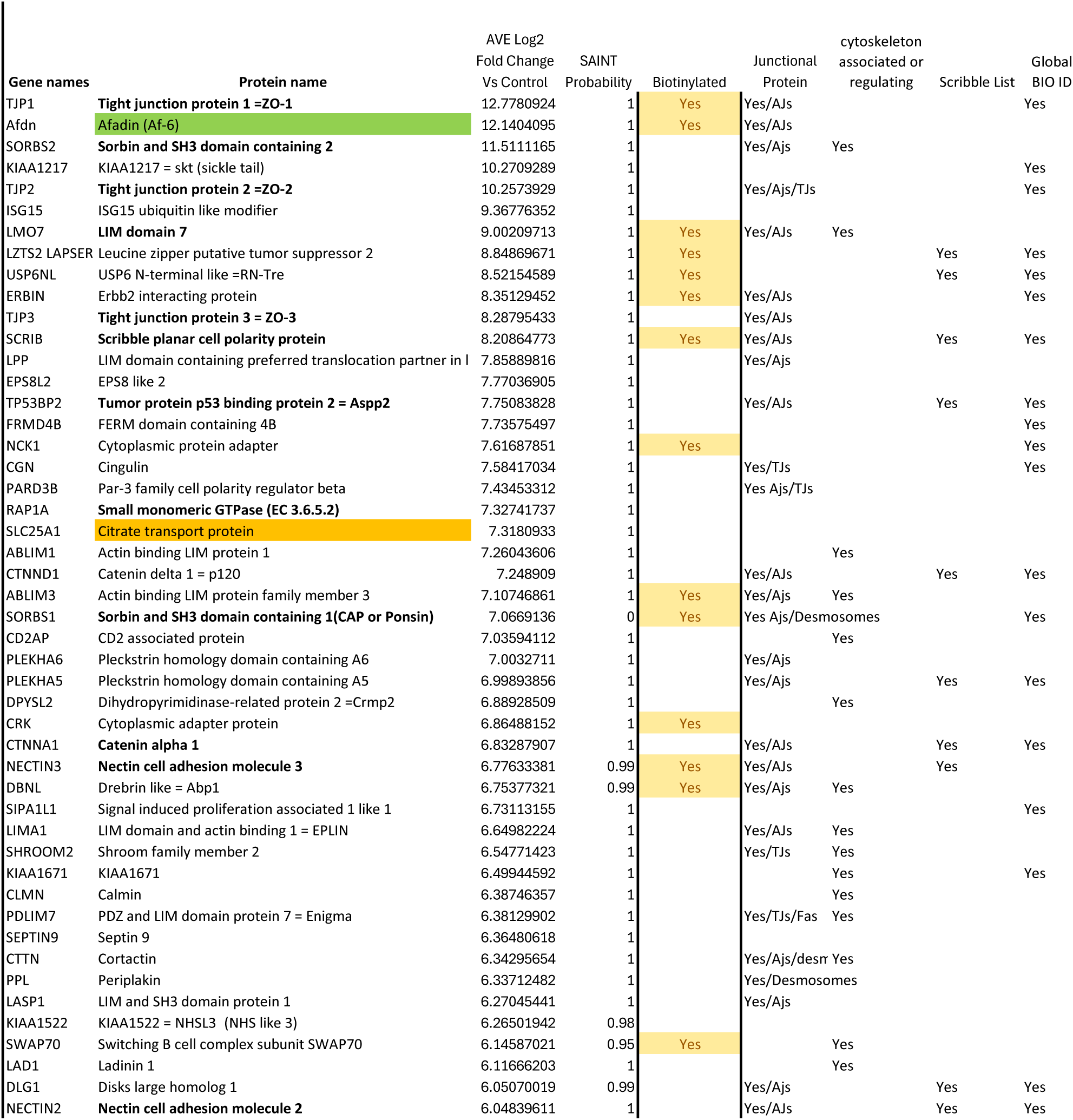
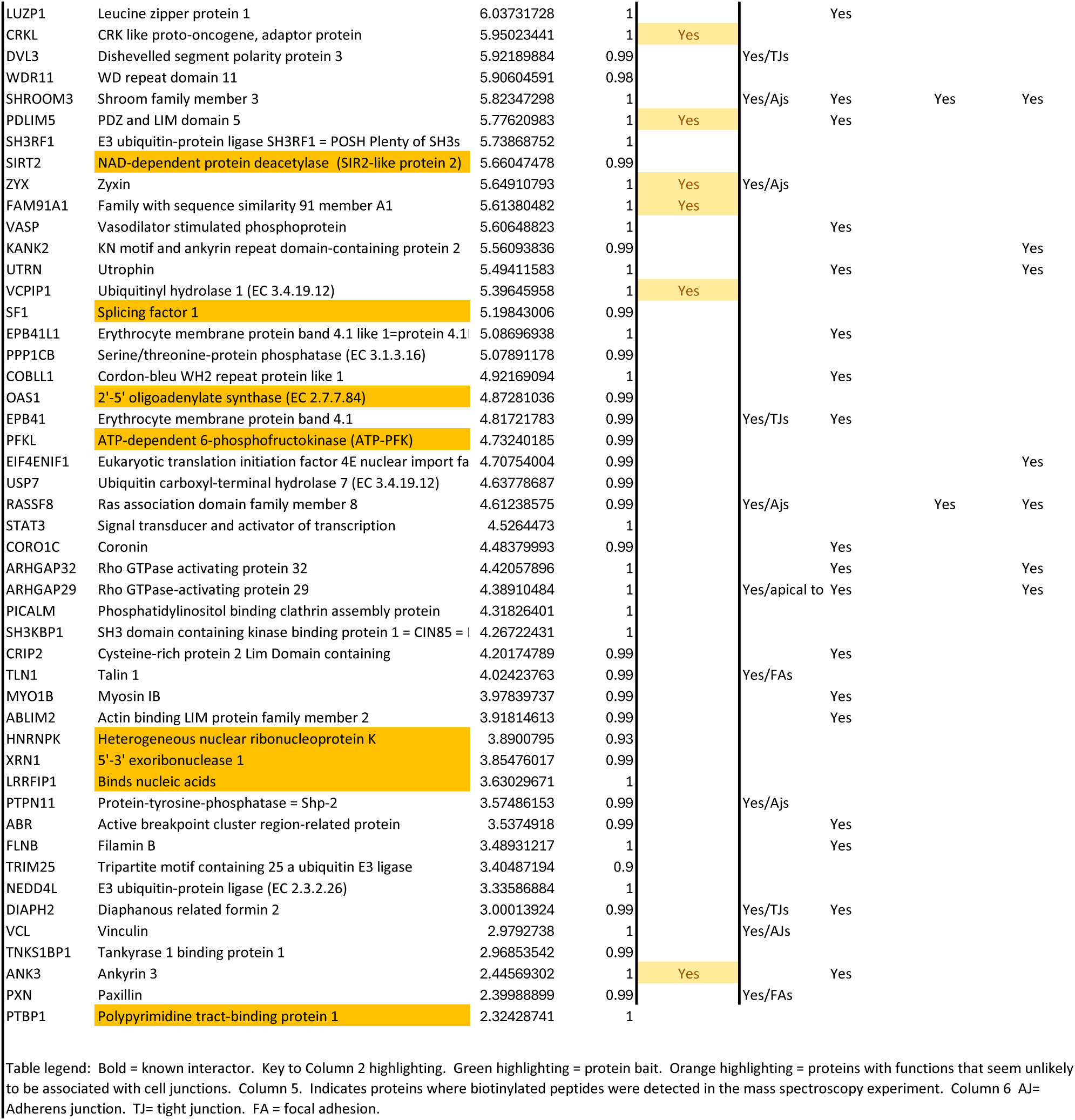
Validated interactors with BL-Afadin.

**Table 2.**
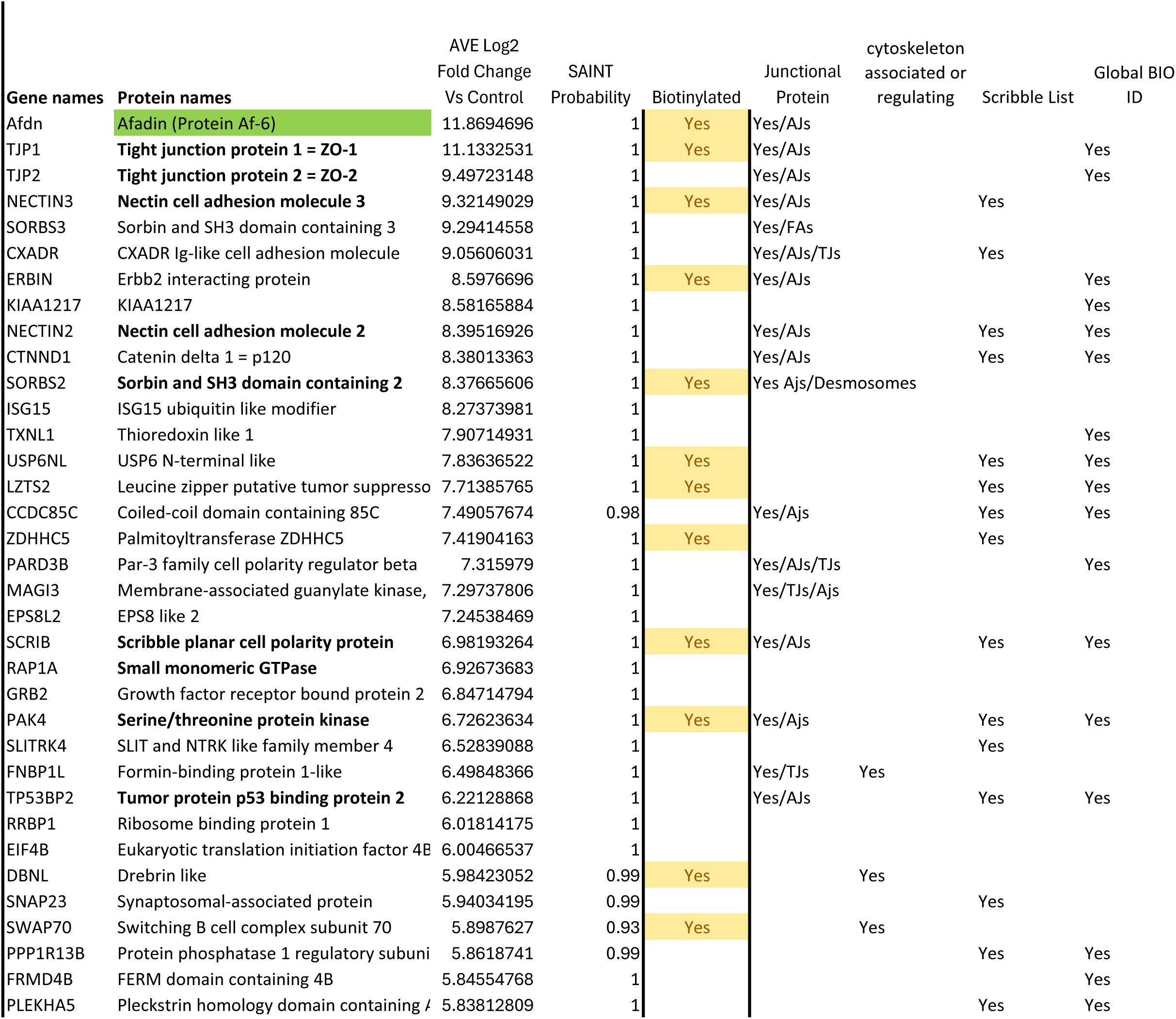
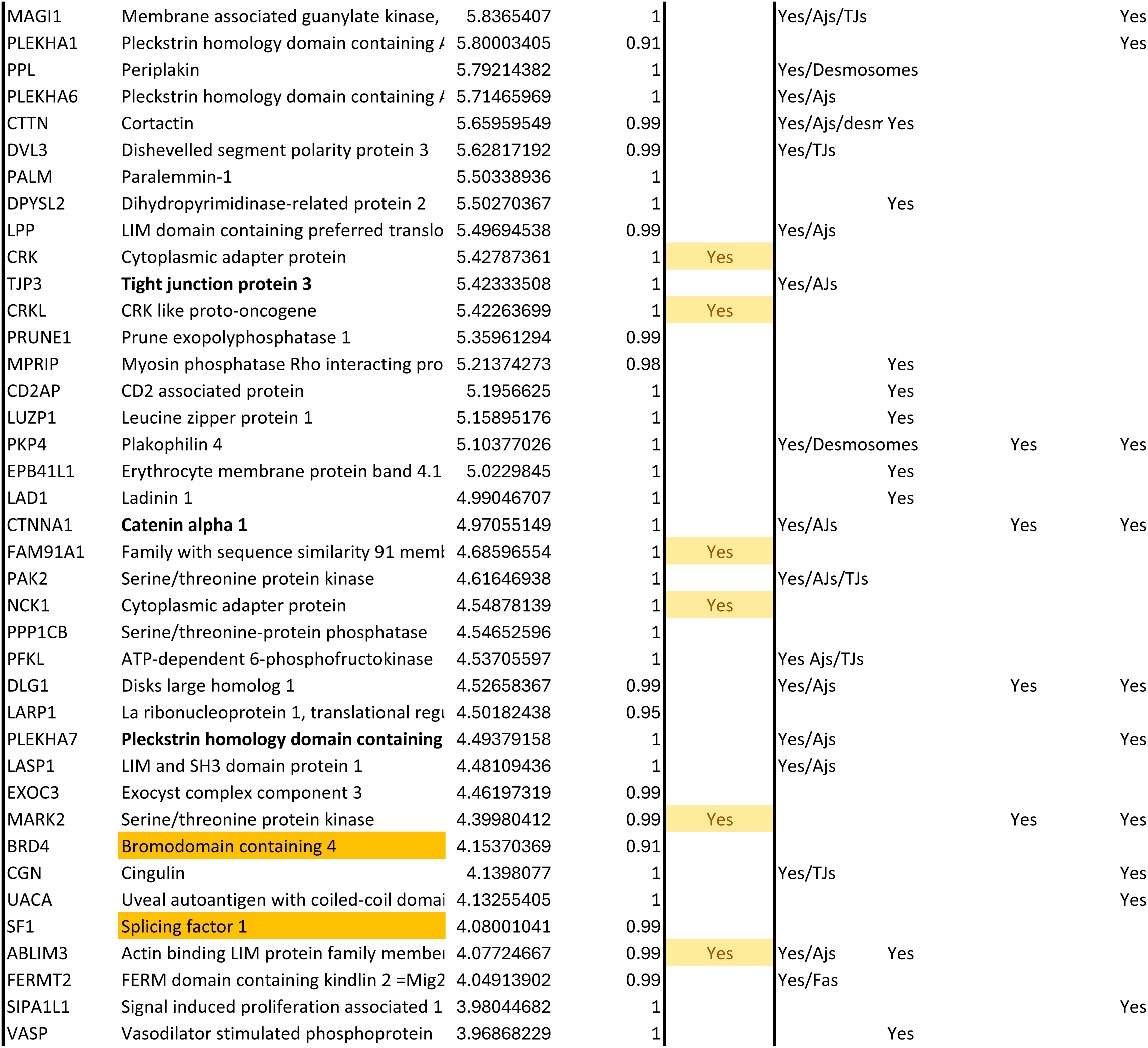
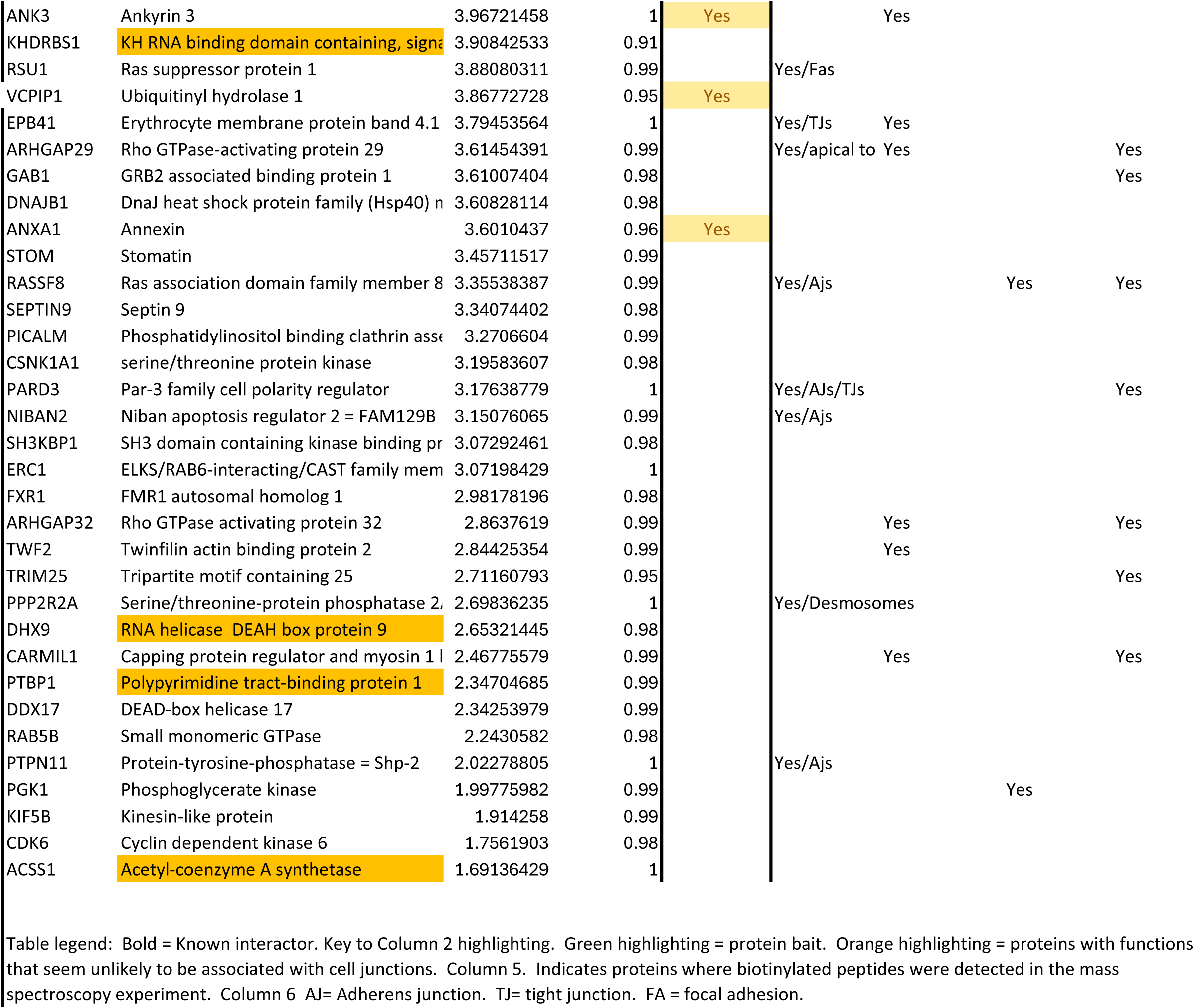
Validated interactors with Afadin-BL.

**Table 3.**
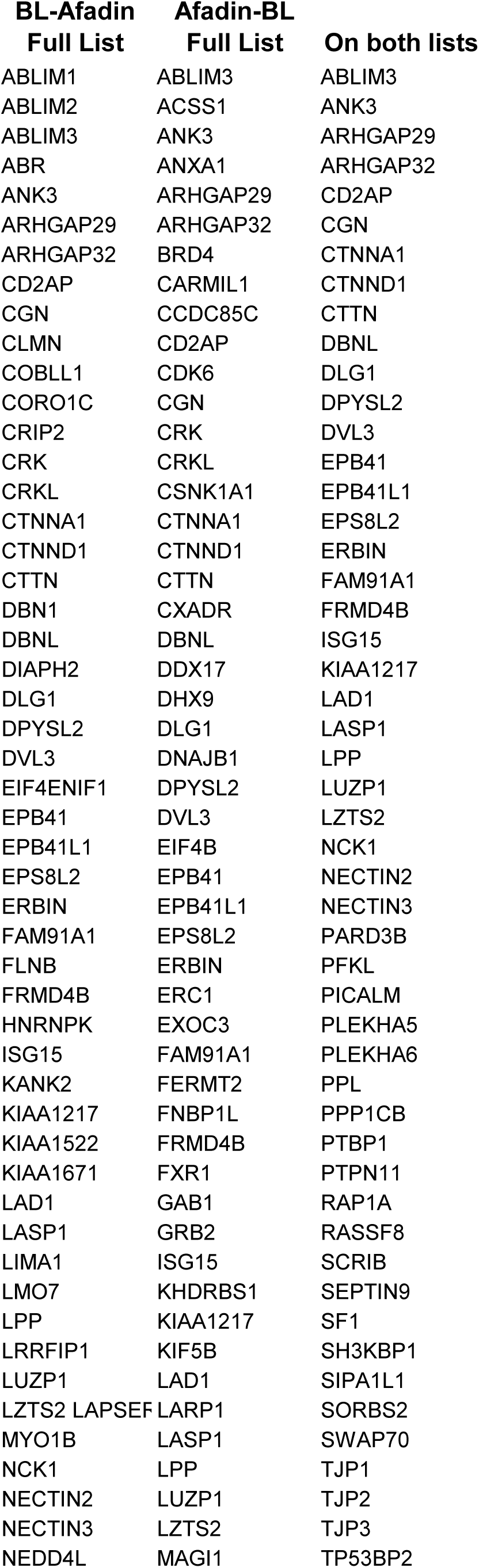
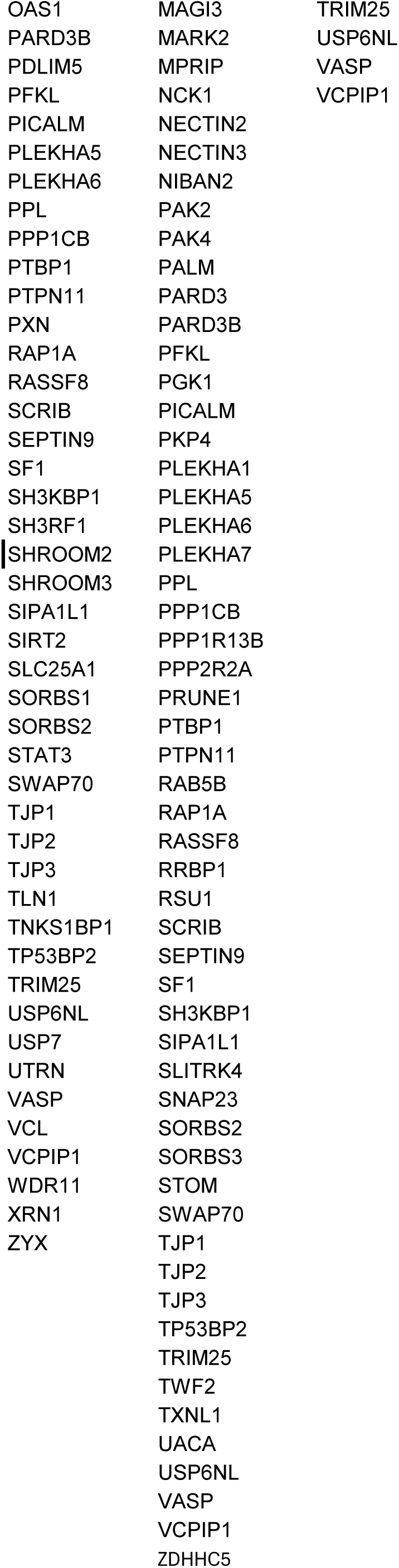
Overlap between the BL-Afadin and Afadin-BL lists (arranged alphabetically)

### Our lists contain many known interactors and are highly enriched for known junctional and cytoskeletal proteins

As a first verification of the quality of our data, we asked whether the lists included proteins known to interact with Afadin directly or via co-immunoprecipitation. The results were striking. Nectins were the first proteins found to interact with Afadin (Takahashi et al., 1999) and both Nectin2 and Nectin3 were on our lists, with SAINT probabilities of 1, 1 or 0.99, 1 for N- or C terminal tagged Afadin respectively. The known binding partner Alpha-catenin (CTNNA1; Pokutta et al., 2002) was also on both N- and C-terminal lists, as was another component of the cadherin-catenin complex, p120 (CTNND1). All three ZO-1 family members (TJP1 (Yamamoto et al., 1997), TJP2, TJP3) were on both lists, as was Rap1A, which binds to Afadin’s N-terminal RA domain (Wohlgemuth et al., 2005) and activates Afadin. Two known interactors identified in previous BioID screens, Scribble (Goudreault et al., 2022) and Pak4 (Baskaran et al., 2021), were also on our lists—intriguingly Pak4 was only identified with our C-terminally tagged Afadin. Four other known interactors were also included: TP53BP2/ASPP2 (Royer et al., 2022), identified with both baits, LMO7 (Ooshio et al., 2004), SORBS1/Ponsin (Mandai et al., 1999), all scoring positive only with the N-terminal bait, and PLEKHA7 (Kurita et al., 2013), scoring positive only with the C-terminal bait.

As a more complete assessment of the overlap between our lists and known interactors, we used The Biological General Repository for Interaction Datasets (BioGRID; Oughtred et al., 2021), a public database that archives and disseminates genetic and protein interaction data from model organisms and humans. These are curated from both high-throughput datasets and individual focused studies in the literature. Using the data from humans (the closest match to the canine cells we used), 294 proteins are annotated as interacting with Afadin genetically or via protein-protein interactions (Table S2). Of the 141 proximal proteins identified on one or both of our lists, 43 were previously reported as physically co-complexed with Afadin within the BioGrid database (Oughtred et al., 2021). These include 7 found only on our BL-Afadin list, 16 found only on our Afadin-BL list, and 20 proteins found on both (Fig. 4A; Table S2).

**Fig. 4.**
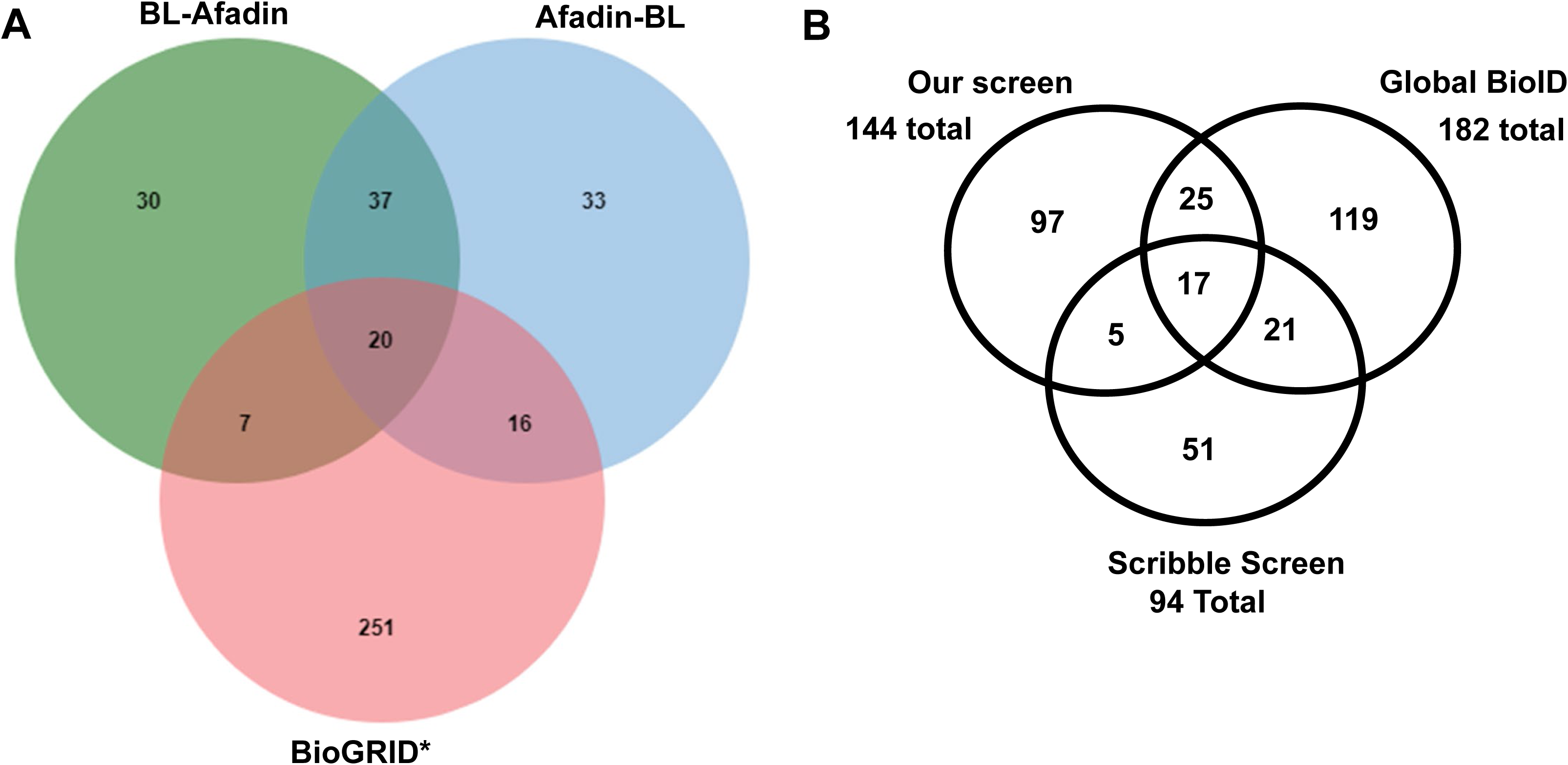
Venn diagrams illustrating the overlap between different screens. A. Overlap between the full list of proteins in the BioGRID database as interacting with Afadin, and the lists of proteins identified as Afadin neighbors using BL-Afadin or Afadin-BL. B. Overlap of the proteins identified in our screen with both baits and those identified in two earlier BioID screens with N-terminally tagged Afadin.

Next, we used the Panther 19.0 classification system (Thomas et al., 2022) to determine the Gene Ontology (GO) biological processes over-represented in our lists relative to the full set of 20580 proteins in the human genome. We used proteins with a SAINT score = 1 from the BL-Afadin or Afadin-BL lists. Each list was enriched for similar but not identical biological processes. The most enriched process on the BL-Afadin list was “cytoskeletal organization”, followed by “actin-filament-based process and “actin cytoskeleton organization” (Table S3). “Cell adhesion” was the fifth most enriched process, and “cell junction organization” and “cell-cell adhesion” were in the top ten, along with two processes involved in “barrier” function. The most enriched process on the Afadin-BL list was “cell-cell junction organization”, with “cell junction organization”, “cell adhesion” “cell-cell adhesion”, and “protein localization to cell-cell junction” ranked third, fourth, fifth, and sixth, while two processes involved in “barrier” function were in the top ten (Table S4). “Cytoskeletal organization” was ranked second. Thus, our lists are statistically strongly enriched for cell junction and cytoskeletal proteins.

We also manually examined the literature to examine what was known about the subcellular localization of the hits with SAINT scores ≥ 0.9 on the BL-Afadin, Afadin-BL, or both lists (Tables 1 and 2; total = 146 proteins on one, the other, or both lists). 39 proteins are reported to localize to AJs, and an additional 14 are known to localize to tight junctions (some are reported to localize to both). Intriguingly, our lists also included 5 proteins reported to localize to desmosomes, including two well-known desmosomal adapter proteins, Periplakin and Plakophilin4. The lists also include many proteins known to regulate or associate with the actomyosin cytoskeleton—39 proteins fit that category.

### Afadin proximal proteins in ZO-knockdown MDCK cells

One of the secondary aims of our experiments was to compare the protein neighbors of Afadin in wildtype MDCK cells and in MDCK cells depleted for the tight junction proteins of the ZO-1 family (ZO-KD cells). In our previous work we found that shRNAi-based knockdown of ZO-1 and ZO-2 dramatically altered junctional architecture (Choi et al., 2016; Fanning et al., 2012). Cell junctions became much straighter, and robust sarcomeric arrays of actin and myosin assembled along AJs. In response to the elevated junctional tension, Afadin recruitment to junctions was elevated and it was important for maintaining epithelial architecture in these cells. We thus wondered if the protein neighbors of Afadin would be altered, so we compared differential enrichment of proteins in wildtype MDCK cells versus ZO-KD cells. One limitation of this analysis was that we only had a single replicate of the ZO-KD samples, and thus combined the BL-Afadin and Afadin-BL samples in this comparison. This also meant fewer proteins met the SAINT ≤0.9 threshold (Table S1). 30 of the 41 proteins on this list were shared with either the BL-Afadin or Afadin-BL lists.

In examining differential expression, we were surprised to find relatively few differences. We did observe the expected reduction of ZO-1 and ZO-2, the proteins that were knocked down—they were down 5.3 and 7.6-fold. ZO-3 was also reduced, consistent with the fact that ZO-3 is less stable when the other family members are knocked down (Fanning et al., 2012). However, the list of proteins relatively enriched in ZO-KD cells was not particularly informative. The three most enriched were either proteins about which little is known (ERICH6B) or enzymes with no clear connection to junctions (acetyl-CoA carboxylase or a component of the pyruvate dehydrogenase complex). One interesting upregulated protein was Vinculin, which is known to be recruited to AJs under elevated tension and which we found enriched at AJs after ZO-KD (Choi et al., 2016). However, in our current dataset this did not get a SAINT score of ≥0.9. The results were similar when we examined proteins differentially enriched in wildtype MDCK cells. Two of the most upregulated proteins seemed to have little connection to cell junctions or the cytoskeleton: the ISG15 ubiquitin like protein ISG15 and the translational regulator LARP1. However, others were more intriguing, including Zyxin (up 4.2 fold) and VASP (up 3.5 fold), which are both recruited to actin filaments after stress or damage (Smith et al., 2010), or MAGI3 (up 4.0 fold), the homolog of which regulates AJs and cell shape changes in the Drosophila eye (Zaessinger et al., 2015). Following up on these differences may help explain the dramatic differences in junctional architecture in these two cell lines.

### Comparing our screen to previous proximity labeling screens

Two previous papers reported the results of proximity labeling screens using Afadin as a bait. Both used a single bait, with Afadin N-terminally tagged. We compared our results to theirs. The first screen was part of a large-scale proteomics effort in which more than 200 intracellular proteins from 32 different cellular compartments were tagged with BirA* and expressed in HEK293 cells (Go et al., 2021). N-terminally tagged Afadin was one of their baits. 26 of the 95 hits from our screen with BL-Afadin with a SAINT score ≥0.9 were also on their list (Table 1) and 34 of the 106 hits from our screen with Afadin-BL with a SAINT score ≥0.9 from our screen were also on their list (Table 2), for a total of 42 shared proteins (Fig. 4B; Table S5), revealing strong overlap. This direct comparison likely understates the overlap, as in many cases one list includes paralogs of proteins on the other list: e.g., Nectin 2 versus Nectin 3 or SorbS1 versus SorbS2. These would add 13 more matches. A second group was interested in Afadin’s role as a Ras GTPase effector and carried out a proximity proteomics screen using N-terminally tagged Afadin as a bait in HeLa cells (Goudreault et al., 2022). Here overlap was present but less pronounced. Only 12 of 95 hits from our screen with BL-Afadin with a SAINT score ≥0.9 (Table 1), and 21 of 106 hits from our screen with Afadin-BL with a SAINT score ≥0.9 (Table 2) were also on their list, for a total of 22 shared proteins (Fig. 4B; Table S5).

Intriguingly, each screen lacked some proteins seen in other screens. Only 17 proteins were found in all three screens (Table S5), and this list only included 5 of the 14 known interactors identified in our screen, lacking for example ZO-1 family proteins. Among the proteins shared by our list and the Global list, there were 43 proteins shared by both of these screens, and the shared set now included 8 of 14 known interactors identified on our lists. As we discuss below, the three screens used three different cell types, likely explaining some of these differences.

## Discussion

Afadin and its Drosophila homolog Canoe are key components of the protein network that links AJs to the actomyosin cytoskeleton. They stabilize this connection as force is generated, thus helping ensure cell shape change and movement during embryonic morphogenesis, organogenesis and tissue homeostasis, without tissue disruption. While initial work suggested E-cadherin links to actin via a simple direct connection involving beta- and alpha-catenin, we now know this connection involves many more proteins linked by multivalent interactions. Many proteins in this network have been defined, but others are likely to be involved. Proximity proteomics provides a powerful tool to identify potential new players.

### Multiple proximity proteomics screens identify overlapping but not identical proteins

Our screen was the third to use N-terminally tagged Afadin as a bait. Several criteria suggest our screen was of high quality. First, we identified 14 known interacting proteins, including the many known physical interactors identified by direct protein interactions or co-immunoprecipitation, along with 43 proteins on the Biogrid interactor list (Table S2). It is possible some known interactors that were not identified, like Nectin1, are not expressed at high enough levels in MDCK cells. Second, when we used the Panther 19.0 classification system to assess the biological processes in which our identified proteins participated, the top categories almost all reflected cell junctions or the cytoskeleton (Tables S3 and S4).

Each of the three screens identified overlapping sets of proteins, with each also including some unique potential protein neighbors. The overlap with one of the previous screens was particularly strong, with 42 out of 146 proteins on at least one of our lists (BL-Afadin or Afadin-BL SAINT ≥ 0.9) shared with that screen (Fig. 4B; Table S5). However, list of proteins overlapping in all three screens (17) or in the two most extensive screens (42), and the fact that neither of these overlapping lists included all known interactors found in our screen, suggests the combined lists are a more useful resource than any single list alone. Another value of the multiple screens is that they targeted different cell types (MDCK cells (our screen), HeLa cells (Goudreault et al., 2022), HEK293 cells (Go et al., 2021)). This helps identify tissue- or cell type-specific interactors. It’s worth noting that HeLa cells, a cervical cancer cell line, do not form fully functional tight junctions and therefore lack robust epithelial barrier function (Shi et al., 2020). Likewise, HEK293 cells do not form tight junctions and are often used as a “tight junction free: background in which to test the function of tight junction proteins like claudins (Inai et al., 2009). The absence of tight junctions seems likely to alter the molecular architecture of other cell junctions. Other proteomics approaches also will provide insights. Afadin was one of the proteins analyzed in a large scale IP-mass spectroscopy screen (Huttlin et al., 2021)—of the 22 proteins identified by IP-mass spectroscopy, eight were shared with our lists (Table S5), but this screen also identified additional proteins we did not identify, and only one of the proteins common between our screen and the IP-mass spectroscopy screen, CXADR, was found in the overlap between all three different proximity proteomics screens.

### How did using both N-terminal and C-terminal baits inform our understanding?

The use of bait proteins tagged with BirA* in different positions has the theoretical advantage of identifying proteins interacting with different protein domains positioned along the N-to C-terminal axis, or that are differentially positioned in a multiprotein complex. Afadin is a relatively large protein (1882 amino acids) with five folded protein domains and a long C-terminal intrinsically disordered region (IDR), and thus the potential distance between an N- and C-terminal BirA* tag is substantial. The 800 aa IDR alone could reach ∼280 nm if fully extended and the theoretical labeling radius of BirA* is ∼10-50 nm (Bosch et al., 2021). However, we did not see dramatic differences between the proteins identified with our two baits. 58 of 145 proteins with SAINT scores ≥ 0.9 were present on <u>both</u> lists. 38 only passed that SAINT score threshold on the BL-Afadin list, and 49 only passed that SAINT score threshold on the Afadin-BL list. We also analyzed overall differential enrichment of all of the proteins we identified (Table S1—Differential enrichment). These differentially enriched proteins may provide clues as to positioning relative to different domains or regions of Afadin—for example, both Nectins we identified, which interact with the PDZ domain, were only found in the Afadin-BL sample, consistent with the idea that the PDZ is the most C-terminal of the folded domains and the IDR may be very flexible. It will be interesting to determine the nature of the interaction between Pak4 and Afadin, as it was the protein most differentially enriched in the Afadin-BL sample (6 fold; Table S1—Differential enrichment). In contrast, three of the LIM domain proteins in the dataset, LMO7, Zyxin, and PDLIM7, were all differentially enriched in the BL -Afadin sample (3.8-4.8 fold; Table S1—Differential enrichment). Their sites of interaction with Afadin remain unknown. At least one differentially-enriched protein, ZO-3, was puzzling given what we know. ZO-1 family proteins can interact with a proline-rich region in the IDR (Ooshio et al., 2010), but ZO-3 was differentially enriched in the BL-Afadin sample (3.5 fold; Table S1—Differential enrichment), which does not fit with this binding site. However, neither of the other ZO-1 family proteins met the threshold of 2-fold differential enrichment.

Several things may explain the strong overlap of proteins identified with the two baits. First, if the IDR is as flexible and extendable as predicted, the C-terminal BirA* may be able to reach proteins throughout the neighborhood. Second, it has become increasingly clear that connections between the cadherin tail and actin do not occur through simple linear connections, as once thought, but involve a large network of proteins linked by multivalent connections. Further, many of the proteins in the linkage network can phase separate (Sun et al., 2022), including Afadin (Kuno et al., 2024), and junctional puncta contain hundreds to thousands of junctional proteins (McGill et al., 2009). Within these potential biomolecular condensates, each Afadin may be in a somewhat different environment, with different binding partners and different neighbors. This contrasts with a model in which each individual Afadin is bound to all of its known binding partners simultaneously, with these fixed interactions creating a more “crystalline” interaction network. In fact, when examining a protein with a long IDR, different parts of the same protein may localize differently—this was observed when using superresolution microscopy to localize N-versus C-terminally tagged ZO-1 (Nguyen et al., 2024; Spadaro et al., 2017).

### Proteomics suggest proteins from “different cell junctions” can be juxtaposed

Textbooks depict the lateral borders of epithelial cells as lined by discrete sets of cell junctions. Apical tight junctions, their more basal neighbors the AJs and the desmosomes found in some epithelial cells, were initially defined by electron microscopy, with cellular and molecular studies adding lists of core components and associated proteins. Recent work added more nuance to this, with high resolution microscopy placing the Crumbs complex even more apical in the “marginal zone” (Martin et al., 2021) and most recently the separation of the AJ into zones enriched for Afadin/Nectins and the cadherin-catenin complex (Mangeol, 2024).

Proximity proteomics provides a tool to explore the distinctions between these junctions and offered some surprises. Van Itallie et al. used proximity proteomics to explore proteins in the vicinity of the core AJ protein E-cadherin (Van Itallie et al., 2014) or in the vicinity of the protein often used as the tight junction marker, ZO-1 (Van Itallie et al., 2013). When we examined the top 125 proteins in each screen, we were surprised to find more matches in the ZO-1 dataset (37 matches; Afadin ranked 15th) than in the E-cadherin dataset (29 matches; Afadin ranked 17th). Tan et al. used proximity proteomics to explore neighbors of polarity regulators in the Par complex—here they used Par3, usually viewed as a tight junction protein, as bait—or of the Crumbs complex, which defines the marginal zone, using Pals as bait (Tan et al., 2020). Consistent with the overlap of our list with that for ZO-1, 18 of 87 proteins on the Par3 list overlapped our lists, including Afadin itself, Nectin2, and alpha-catenin, all usually viewed as AJ proteins. While the Pals list did not include Afadin, it did include Nectin2, and 11 of 110 proteins on this list overlapped with ours, including the tight junction proteins ZO-1, ZO-3, Cingulin.

This suggests additional complexity. First, at the boundaries between different junctions, proteins will likely be close to proteins in the next junction more apical or basal. Recent lovely work from the Honigman lab revealed that interactions between the tight junction protein ZO-1 and the marginal zone protein Patj are key for positioning the tight junction belt apically (Pombo-Garcia et al., 2024). It will be interesting to see if similar mechanisms underlie the recent observation that in at least some cell types Afadin and the Nectins segregate apically to the cadherin-catenin complex (Mangeol, 2024). Second, the neighbors of a protein may change as junctions form and mature in a single cell type—proximity proteomics of ZO-1 supports this idea (Pombo-Garcia et al., 2024). Third, many proteins may localize to more than one junctional type, with the difference being one of degree of enrichment. This may help explain the inclusion on our list of proteins thought to be solely components of desmosomes or integrin-based focal adhesions. These include both proteins known to have a dual localization to different junctions like Plakophilin4 (desmosomes and AJs) or vinculin (focal adhesions or AJs), and proteins usually thought to be only found in desmosomes, like Periplakin, or only in integrin-based junctions, like Talin.

### Proximity proteomics opens new questions and research directions

Proximity proteomics experiments identify potential new partners of the protein of interest and offer leads for new research. The Afadin screens offer three examples. Two groups, including ours, used identification of the polarity regulator Scribble as an Afadin interactor to explore new roles for both proteins: Scribble as a key player in localizing apical AJs during Drosophila development (Bonello et al., 2019) and Afadin as a coupler of Ras GTPases to Scribble to regulate cell polarity and migration (Goudreault et al., 2022). Others used the Global BioID Afadin list to identify the kinase Pak4 as a regulator of junctional protein phosphorylation (Baskaran et al., 2021). Similarly, the ZO-1 BioID screen stimulated studies revealing that the BAR-domain protein TOCA-1 regulates actin assembly at tight junctions (Van Itallie et al., 2015). The new lists offer many exiting new leads to follow. Even if we confine our interests to genes shared in all three Afadin screens, there are intriguing leads. For example, the Rab5-GAP US6NL/RN-Tre regulates integrin endocytosis and focal contact turnover in mammalian cells (Palamidessi et al., 2013) and regulates assembly of myosin into contractile networks in Drosophila cells (Platenkamp et al., 2020)—exploring its role in cell-cell AJs seems well worth the effort. The orthologs of RASSF8 and ASPP1/ PPP1R13B form a complex regulating cell-cell adhesion during Drosophila retinal morphogenesis (Langton et al., 2009), working together with MAGI (Zaessinger et al., 2015), another hit in two of the screens. Looking more broadly at their roles in junctional remodeling during embryonic morphogenesis seems warranted. Little is known about the normal physiological roles of LZTS2 in either mammals or Drosophila—this might also be a fruitful area to pursue. We hope this dataset will stimulate further research into the roles of many of the identified proteins in cell-cell junctions in both mammals and Drosophila.

## Acknowledgements

We are very grateful to Tina van Italie for advice on the use of BioID, and Dhaval Bhatt for assistance with ordering proteins by the difference in average log2 LFQ intensity values between experimental and control groups.

## Methods

### Stable expression of BirA*-Afadin in MDCK cells

We utilized fusion proteins we had previously generated (Bonello et al., 2019). Rat Afadin coding sequences were cloned into an inducible mammalian expression vector for BioID, pTRE2hyg-BirA*-myc (Van Itallie et al., 2013). We created versions with Afadin coding sequences tagged with BirA* at the N-(pTRE2hyg-BirA*-Afadin-myc) or C-terminus (pTRE2hyg-Afadin-BirA*-myc). We then transfected subconfluent MDCK T23 cells with these plasmids plasmids, or a plasmid encoding BirA*-myc alone, and selected for the stable clones using hygromycin as a selective drug. Stable clones were verified by immunoblotting and by and immunostaining for Afadin and the Myc tag carried on the fusion proteins. We cultured cells expressing our BirA* fusions in DMEM media containing 1 g/l glucose, 10% fetal bovine serum, 15 mM HEPES (pH 7.4) and 50 ng/ml doxycycline, to keep transgene expression off.

### Immunostaining (cell culture)

MDCK and ZOdKD (Choi et al., 2016) cells expressing BioID constructs were cultured for 7 days in Transwells without doxycycline to express BirA*-Afadin fusions and fixed in ice-cold ethanol for 1 h at −20°C. After three washes with PBS, the samples were incubated with blocking buffer (10% FBS in PBS) for 1 h at room temperature. Subsequently, the inserts were incubated with primary antibodies diluted in blocking buffer. Rabbit anti-Afadin and mouse anti-Scribble, or rabbit anti-Scribble and mouse anti-Myc antibodies were used for co-staining. Following three washes with wash buffer (1% FBS in PBS), the cells were incubated with the Alexa-conjugated secondary antibodies along with Hoechst 33343 to stain DNA. After three washes with wash buffer, the insert membrane was cropped out, mounted on a microscope slide with Prolong Diamond antifade mountant (Thermo Fisher Scientific) and cured before imaging.

### Expression and purification of biotinylated proteins

MDCK cells were those used by Choi et al. (2016) and were authenticated and tested for contamination. MDCK cells stably expressing BirA* transgenes were cultured without doxycycline for 5 days, following which cells were treated with 50 µM biotin for 24 h. To prepare lysates, cells were washed three times with ice-cold PBS and scraped into lysis buffer [1% NP-40, 0.5% deoxycholate, 0.2% SDS, 50 mM Tris (pH 8), 150 mM NaCl, 2 mM EDTA, supplemented with protease/phosphatase inhibitor]. Lysates were snap frozen on dry ice prior to storing at −80°C. To capture biotinylated proteins, lysates were thawed at 4°C, sonicated (amplitude 50%, 10 strokes performed manually) and incubated on ice for 15 min. Lysates were then spun at 15,000 ***g*** for 15 min and the protein concentration of supernatant determined using BioRad Protein Assay Dye. Equal concentrations of sample were added to pre-washed Dynabeads (MyOne Streptavidin C1) and incubated with nutation overnight at 4°C. After removing the unbound sample, beads were washed twice with buffer 1 (2% SDS) for 10 min, once with buffer 2 [0.1% deoxycholate, 1% Triton X-100, 500 mM NaCl, 1 mM EDTA and 50 mM Hepes (pH 8)] for 10 min, once with buffer 3 [0.5% deoxycholate, 0.5% NP-40, 250 mM NaCl, 1 mM EDTA and 10 mM Tris (pH 8)] for 10 min and twice with buffer 4 [50 mM NaCl and 50 mM Tris (pH 7.4)] for 10 min. After last wash, the beads were incubated with 5 mM DTT for 15 min at 60C. The beads slurry was applied to a spin filter column (VIVACON 500, 30,000 MWCO, Fisher Scientific) and centrifuged at 10,000 x g for 20 min. The column was washed three times with 8M urea, and incubated in 100 uL of chloroacetamide:urea (1:9) solution for 20 min in the dark, followed by rinse twice with 8M urea. The denatured proteins were stabilized by washing twice with 50 mM ammonium bicarbonate and subjected to trypsin digestion (Promega, V115C) by incubating at 37C overnight. The tryptic peptides were collected by centrifugation at 10,000 x g for 10 min, and the bead-trapped peptides were eluted with a high-temperature, high-organic method (0.5% trifluoroacetic acid/acetonitrile (4:6) at 65C for 30 min). The collected peptides were cleaned with C-18 spin column, vacuum dried and reconstituted in Buffer A [0.1% TFA in ddH2O]. The dissolved peptides were loaded onto the mass spectrometer.

### Mass spectrometry data acquisition

Trypsinized peptides were separated via reverse-phase nano-HPLC using a nanoAquity ultra performance liquid chromatography (UPLC) system (Waters Corp.). Peptides were first trapped in a 2-cm trapping column (Acclaim® PepMap 100, C18 beads of 3.0-μm particle size, 100-Å pore size) and a 25-cm EASY-spray analytical column (75-μm inner diameter, C18 beads of 2.0-μm particle size, 100-Å pore size) at 35 °C. The flow rate was 250 nl/min over a gradient of 5% buffer B (0.1% formic acid in acetonitrile) to 35% buffer B in 150 min, and an in Orbitrap Elite mass spectrometer (Thermo Scientific) performed mass spectral analysis. The ion source was operated at 2.4–2.8 kV with the ion transfer tube temperature set at 300 °C. MS1 spectra (300–2000 *m*/*z*) were acquired by the Orbitrap analyzer with 120,000 resolution. MS2 spectra were acquired in data-dependent mode on the 15 most intense peaks by the linear ion trap. The MS2 isolation window as 2.0 *m/z* wide and the normalized collision energy was 35%. The precursor ions were selected based on charge states (+2) and intensity thresholds (above 1e5) from the MS1 scan; dynamic exclusion (one repeat during 30 s, a 60-s exclusion time window, 15ppm tolerance) was also used. The mass spectrometry proteomics data have been deposited to the ProteomeXchange Consortium via the PRIDE partner repository with the dataset identifier PXD056971.

### Mass spectrometry data processing

Raw MS data files were processed by MaxQuant (version 2.4.2.0; (Tyanova et al., 2016)) using the UniProtKB *Canus lupis familiaris* canonical sequence database (downloaded Dec. 2023) and the human Afadin entry (UniProt Accession O35889; (UniProt, 2019). The following parameters were used: specific tryptic digestion with up to two missed cleavages, fixed carbamidomethyl modification, variable modifications for protein N-terminal acetylation, methionine oxidation, lysine biotinylation, match between runs, and label-free quantification. Prey proteins were filtered for high-confidence physical interactions and proximal proteins by scoring with SAINTexpress (v3.6.3) (Teo et al., 2014). SAINT was executed three separate times to score the following conditions: 1) Afadin-BL vs control, 2) BL-Afadin vs control, and 3) Afadin in ZOdKD cells vs control. A SAINT threshold of AvgP >=0.9 was used in all cases.

### Detailed author contributions

Wangsun Choi, Alan Fanning and Mark Peifer conceived the project with advice from Ben Major and Dennis Goldfarb. Wangsun Choi carried out all of the cell biological work, up to preparing the samples for mass spectroscopy. Feng Yan and Ben Major carried out the mass spectroscopy and Dennis Goldfarb analyzed the mass spectroscopy data. Mark Peifer, Ben Major and Dennis Goldfarb wrote the paper with editorial contributions from the other authors.

## References

Asada, M., Irie, K., Morimoto, K., Yamada, A., Ikeda, W., Takeuchi, M. and Takai, Y. (2003). ADIP, a novel Afadin- and alpha-actinin-binding protein localized at cell-cell adherens junctions. J Biol Chem 278, 4103–4111.

Baskaran, Y., Tay, F. P., Ng, E. Y. W., Swa, C. L. F., Wee, S., Gunaratne, J. and Manser, E. (2021). Proximity proteomics identifies PAK4 as a component of Afadin-Nectin junctions. Nat Commun 12, 5315.

Boettner, B., Govek, E. E., Cross, J. and Van Aelst, L. (2000). The junctional multidomain protein AF-6 is a binding partner of the Rap1A GTPase and associates with the actin cytoskeletal regulator profilin. Proc Natl Acad Sci U S A 97, 9064–9069.

Boettner, B., Harjes, P., Ishimaru, S., Heke, M., Fan, H. Q., Qin, Y., Van Aelst, L. and Gaul, U. (2003). The AF-6 homolog canoe acts as a Rap1 effector during dorsal closure of the Drosophila embryo. Genetics 165, 159–169.

Bonello, T. T., Choi, W. and Peifer, M. (2019). Scribble and Discs-large direct initial assembly and positioning of adherens junctions during the establishment of apical-basal polarity. Development 146.

Bosch, J. A., Chen, C. L. and Perrimon, N. (2021). Proximity-dependent labeling methods for proteomic profiling in living cells: An update. Wiley interdisciplinary reviews. Developmental biology 10, e392.

Buchert, M., Schneider, S., Meskenaite, V., Adams, M. T., Canaani, E., Baechi, T., Moelling, K. and Hovens, C. M. (1999). The junction-associated protein AF-6 interacts and clusters with specific Eph receptor tyrosine kinases at specialized sites of cell-cell contact in the brain. J Cell Biol 144, 361–371.

Buckley, C. D., Tan, J., Anderson, K. L., Hanein, D., Volkmann, N., Weis, W. I., Nelson, W. J. and Dunn, A. R. (2014). Cell adhesion. The minimal cadherin-catenin complex binds to actin filaments under force. Science 346, 1254211.

Calautti, E., Cabodi, S., Stein, P. L., Hatzfeld, M., Kedersha, N. and Paolo Dotto, G. (1998). Tyrosine phosphorylation and src family kinases control keratinocyte cell-cell adhesion. Journal of Cell Biology 141, 1449–1465.

Campas, O., Noordstra, I. and Yap, A. S. (2024). Adherens junctions as molecular regulators of emergent tissue mechanics. Nat Rev Mol Cell Biol 25, 252–269.

Carminati, M., Gallini, S., Pirovano, L., Alfieri, A., Bisi, S. and Mapelli, M. (2016). Concomitant binding of Afadin to LGN and F-actin directs planar spindle orientation. Nat Struct Mol Biol 23, 155–163.

Case, L. B. and Waterman, C. M. (2015). Integration of actin dynamics and cell adhesion by a three-dimensional, mechanosensitive molecular clutch. Nat Cell Biol 17, 955–963.

Choi, W., Acharya, B. R., Peyret, G., Fardin, M. A., Mege, R. M., Ladoux, B., Yap, A. S., Fanning, A. S. and Peifer, M. (2016). Remodeling the zonula adherens in response to tension and the role of afadin in this response. J Cell Biol 213, 243–260.

Choi, W., Harris, N. J., Sumigray, K. D. and Peifer, M. (2013). Rap1 and Canoe/afadin are essential for establishment of apical-basal polarity in the Drosophila embryo. Mol Biol Cell 24, 945–963.

Cronin, N. M. and DeMali, K. A. (2021). Dynamics of the Actin Cytoskeleton at Adhesion Complexes. Biology 11.

Ebnet, K., Schulz, C. U., Meyer Zu Brickwedde, M. K., Pendl, G. G. and Vestweber, D. (2000). Junctional adhesion molecule interacts with the PDZ domain-containing proteins AF-6 and ZO-1. J Biol Chem 275, 27979–27988.

Fanning, A. S., Van Itallie, C. M. and Anderson, J. M. (2012). Zonula occludens-1 and -2 regulate apical cell structure and the zonula adherens cytoskeleton in polarized epithelia. Mol Biol Cell 23, 577–590.

Fedor-Chaiken, M., Hein, P. W., Stewart, J. C., Brackenbury, R. and Kinch, M. S. (2003). E-cadherin binding modulates EGF receptor activation. Cell Commun Adhes 10, 105–118.

Gaengel, K. and Mlodzik, M. (2003). Egfr signaling regulates ommatidial rotation and cell motility in the Drosophila eye via MAPK/Pnt signaling and the Ras effector Canoe/AF6. Development 130, 5413–5423.

Gates, J., Mahaffey, J. P., Rogers, S. L., Emerson, M., Rogers, E. M., Sottile, S. L., Van Vactor, D., Gertler, F. B. and Peifer, M. (2007). Enabled plays key roles in embryonic epithelial morphogenesis in Drosophila. Development 134, 2027–2039.

Gates, J. and Peifer, M. (2005). Can 1000 reviews be wrong? Actin, alpha-Catenin, and adherens junctions. Cell 123, 769–772.

Go, C. D., Knight, J. D. R., Rajasekharan, A., Rathod, B., Hesketh, G. G., Abe, K. T., Youn, J. Y., Samavarchi-Tehrani, P., Zhang, H., Zhu, L. Y., et al. (2021). A proximity-dependent biotinylation map of a human cell. Nature 595, 120–124.

Gong, R., Reynolds, M. J., Sun, X. and Alushin, G. M. (2024). Afadin mediates cadherin-catenin complex clustering on F-actin linked to cooperative binding and filament curvature. bioRxiv.

Goudreault, M., Gagne, V., Jo, C. H., Singh, S., Killoran, R. C., Gingras, A. C. and Smith, M. J. (2022). Afadin couples RAS GTPases to the polarity rheostat Scribble. Nat Commun 13, 4562.

Gurley, N. J., Szymanski, R. A., Dowen, R. H., Butcher, T. A., Ishiyama, N. and Peifer, M. (2023). Exploring the evolution and function of Canoe’s intrinsically disordered region in linking cell-cell junctions to the cytoskeleton during embryonic morphogenesis. PLoS One 18, e0289224.

Higashi, T. and Chiba, H. (2020). Molecular organization, regulation and function of tricellular junctions. Biochim Biophys Acta Biomembr 1862, 183143.

Homem, C. C. and Peifer, M. (2008). Diaphanous regulates myosin and adherens junctions to control cell contractility and protrusive behavior during morphogenesis. Development 135, 1005–1018.

Huttlin, E. L., Bruckner, R. J., Navarrete-Perea, J., Cannon, J. R., Baltier, K., Gebreab, F., Gygi, M. P., Thornock, A., Zarraga, G., Tam, S., et al. (2021). Dual proteome-scale networks reveal cell-specific remodeling of the human interactome. Cell 184, 3022–3040 e3028.

Ikeda, W., Nakanishi, H., Miyoshi, J., Mandai, K., Ishizaki, H., Tanaka, M., Togawa, A., Takahashi, K., Nishioka, H., Yoshida, H., et al. (1999). Afadin: A key molecule essential for structural organization of cell-cell junctions of polarized epithelia during embryogenesis. J Cell Biol 146, 1117–1132.

Inai, T., Sengoku, A., Hirose, E., Iida, H. and Shibata, Y. (2009). Freeze-fracture electron microscopic study of tight junction strands in HEK293 cells and MDCK II cells expressing claudin-1 mutants in the second extracellular loop. Histochem Cell Biol 131, 681–690.

Kovacs, E. M., Goodwin, M., Ali, R. G., Paterson, A. D. and Yap, A. S. (2002). Cadherin-directed actin assembly: E-cadherin physically associates with the Arp2/3 complex to direct actin assembly in nascent adhesive contacts. Curr Biol 12, 379–382.

Kuno, S., Nakamura, R., Otani, T. and Togashi, H. (2024). Multivalent Afadin Interaction Promotes IDR-Mediated Condensate Formation and Junctional Separation of Epithelial Cells. bioRxiv 2024.04.26.591237.

Kurita, S., Yamada, T., Rikitsu, E., Ikeda, W. and Takai, Y. (2013). Binding between the junctional proteins afadin and PLEKHA7 and implication in the formation of adherens junction in epithelial cells. J Biol Chem 288, 29356–29368.

Langton, P. F., Colombani, J., Chan, E. H., Wepf, A., Gstaiger, M. and Tapon, N. (2009). The dASPP-dRASSF8 complex regulates cell-cell adhesion during Drosophila retinal morphogenesis. Curr Biol 19, 1969–1978.

Mandai, K., Nakanishi, H., Satoh, A., Obaishi, H., Wada, M., Nishioka, H., Itoh, M., Mizoguchi, A., Aoki, T., Fujimoto, T., et al. (1997). Afadin: A novel actin filament-binding protein with one PDZ domain localized at cadherin-based cell-to-cell adherens junction. J Cell Biol 139, 517–528.

Mandai, K., Nakanishi, H., Satoh, A., Takahashi, K., Satoh, K., Nishioka, H., Mizoguchi, A. and Takai, Y. (1999). Ponsin/SH3P12: an l-afadin- and vinculin-binding protein localized at cell-cell and cell-matrix adherens junctions. J Cell Biol 144, 1001–1017.

Mangeol, P., Massey-Harroche, D., Sebbagh, M., Richard, F., Le Bivic, A., Lenne. P.F. (2024). The zonula adherens matura redefines the apical junction of intestinal epithelia. Proc. Nat. Acad. Sci. USA in press.

Martin, E., Girardello, R., Dittmar, G. and Ludwig, A. (2021). New insights into the organization and regulation of the apical polarity network in mammalian epithelial cells. FEBS J 288, 7073–7095.

McGill, M. A., McKinley, R. F. and Harris, T. J. (2009). Independent cadherin-catenin and Bazooka clusters interact to assemble adherens junctions. J Cell Biol 185, 787–796.

McParland, E. D., Gurley, N. J., Wolfsberg, L. R., Butcher, T. A., Bhattarai, A., Jensen, C. C., Johnson, R. I., Slep, K. C. and Peifer, M. (2024). The dual Ras Association (RA) Domains of Drosophila Canoe have differential roles in linking cell junctions to the cytoskeleton during morphogenesis. Journal of Cell Science in press.

Miyamoto, H., Nihonmatsu, I., Kondo, S., Ueda, R., Togashi, S., Hirata, K., Ikegami, Y. and Yamamoto, D. (1995). *canoe* encodes a novel protein containing a GLGF/DHR motif and functions with *Notch* and *scabrous* in common developmental pathways in *Drosophila*. Genes Dev. 9, 612–625.

Nguyen, T. P., Otani, T., Tsutsumi, M., Kinoshita, N., Fujiwara, S., Nemoto, T., Fujimori, T. and Furuse, M. (2024). Tight junction membrane proteins regulate the mechanical resistance of the apical junctional complex. J Cell Biol 223.

Ooshio, T., Irie, K., Morimoto, K., Fukuhara, A., Imai, T. and Takai, Y. (2004). Involvement of LMO7 in the association of two cell-cell adhesion molecules, nectin and E-cadherin, through afadin and alpha-actinin in epithelial cells. J Biol Chem 279, 31365–31373.

Ooshio, T., Kobayashi, R., Ikeda, W., Miyata, M., Fukumoto, Y., Matsuzawa, N., Ogita, H. and Takai, Y. (2010). Involvement of the interaction of afadin with ZO-1 in the formation of tight junctions in Madin-Darby canine kidney cells. J Biol Chem 285, 5003–5012.

Oughtred, R., Rust, J., Chang, C., Breitkreutz, B. J., Stark, C., Willems, A., Boucher, L., Leung, G., Kolas, N., Zhang, F., et al. (2021). The BioGRID database: A comprehensive biomedical resource of curated protein, genetic, and chemical interactions. Protein Sci 30, 187–200.

Palamidessi, A., Frittoli, E., Ducano, N., Offenhauser, N., Sigismund, S., Kajiho, H., Parazzoli, D., Oldani, A., Gobbi, M., Serini, G., et al. (2013). The GTPase-activating protein RN-tre controls focal adhesion turnover and cell migration. Curr Biol 23, 2355–2364.

Perez-Vale, K. Z. and Peifer, M. (2020). Orchestrating morphogenesis: building the body plan by cell shape changes and movements. Development 147, dev191049.

Platenkamp, A., Detmar, E., Sepulveda, L., Ritz, A., Rogers, S. L. and Applewhite, D. A. (2020). The Drosophila melanogaster Rab GAP RN-tre cross-talks with the Rho1 signaling pathway to regulate nonmuscle myosin II localization and function. Mol Biol Cell 31, 2379–2397.

Pokutta, S., Drees, F., Takai, Y., Nelson, W. J. and Weis, W. I. (2002). Biochemical and structural definition of the l-afadin- and actin-binding sites of alpha-catenin. J Biol Chem 277, 18868–18874.

Pombo-Garcia, K., Adame-Arana, O., Martin-Lemaitre, C., Julicher, F. and Honigmann, A. (2024). Membrane prewetting by condensates promotes tight-junction belt formation. Nature 632, 647–655.

Popovic, M., Bella, J., Zlatev, V., Hodnik, V., Anderluh, G., Barlow, P. N., Pintar, A. and Pongor, S. (2011). The interaction of Jagged-1 cytoplasmic tail with afadin PDZ domain is local, folding-independent, and tuned by phosphorylation. Journal of molecular recognition : JMR 24, 245–253.

Radziwill, G., Erdmann, R. A., Margelisch, U. and Moelling, K. (2003). The Bcr kinase downregulates Ras signaling by phosphorylating AF-6 and binding to its PDZ domain. Mol Cell Biol 23, 4663–4672.

Rouaud, F., Sluysmans, S., Flinois, A., Shah, J., Vasileva, E. and Citi, S. (2020). Scaffolding proteins of vertebrate apical junctions: structure, functions and biophysics. Biochim Biophys Acta Biomembr 1862, 183399.

Roux, K. J., Kim, D. I., Burke, B. and May, D. G. (2018). BioID: A Screen for Protein-Protein Interactions. Curr Protoc Protein Sci 91, 19 23 11–19 23 15.

Royer, C., Sandham, E., Slee, E., Schneider, F., Lagerholm, C. B., Godwin, J., Veits, N., Hathrell, H., Zhou, F., Leonavicius, K., et al. (2022). ASPP2 maintains the integrity of mechanically stressed pseudostratified epithelia during morphogenesis. Nat Commun 13, 941.

Sawyer, J. K., Choi, W., Jung, K. C., He, L., Harris, N. J. and Peifer, M. (2011). A contractile actomyosin network linked to adherens junctions by Canoe/afadin helps drive convergent extension. Mol Biol Cell 22, 2491–2508.

Sawyer, J. K., Harris, N. J., Slep, K. C., Gaul, U. and Peifer, M. (2009). The Drosophila afadin homologue Canoe regulates linkage of the actin cytoskeleton to adherens junctions during apical constriction. J Cell Biol 186, 57–73.

Sears, R. M., May, D. G. and Roux, K. J. (2019). BioID as a Tool for Protein-Proximity Labeling in Living Cells. Methods Mol Biol 2012, 299–313.

Shi, Y., Li, R., Yang, J. and Li, X. (2020). No tight junctions in tight junction protein-1 expressing HeLa and fibroblast cells. Int J Physiol Pathophysiol Pharmacol 12, 70–78.

Smith, M. A., Blankman, E., Gardel, M. L., Luettjohann, L., Waterman, C. M. and Beckerle, M. C. (2010). A zyxin-mediated mechanism for actin stress fiber maintenance and repair. Dev Cell 19, 365–376.

Smith, M. J. (2023). Defining bone fide effectors of RAS GTPases. Bioessays 45, e2300088.

Spadaro, D., Le, S., Laroche, T., Mean, I., Jond, L., Yan, J. and Citi, S. (2017). Tension-Dependent Stretching Activates ZO-1 to Control the Junctional Localization of Its Interactors. Curr Biol 27, 3783–3795 e3788.

Stevens, T. L., Rogers, E. M., Koontz, L. M., Fox, D. T., Homem, C. C., Nowotarski, S. H., Artabazon, N. B. and Peifer, M. (2008). Using Bcr-Abl to examine mechanisms by which abl kinase regulates morphogenesis in Drosophila. Mol Biol Cell 19, 378–393.

Sun, D., LuValle-Burke, I., Pombo-Garcia, K. and Honigmann, A. (2022). Biomolecular condensates in epithelial junctions. Curr Opin Cell Biol 77, 102089.

Takahashi, K., Matsuo, T., Katsube, T., Ueda, R. and Yamamoto, D. (1998). Direct binding between two PDZ domain proteins Canoe and ZO-1 and their roles in regulation of the jun N-terminal kinase pathway in Drosophila morphogenesis. Mech. Dev. 78, 97–111.

Takahashi, K., Nakanishi, H., Miyahara, M., Mandai, K., Satoh, K., Satoh, A., Nishioka, H., Aoki, J., Nomoto, A., Mizoguchi, A., et al. (1999). Nectin/PRR: an immunoglobulin-like cell adhesion molecule recruited to cadherin-based adherens junctions through interaction with Afadin, a PDZ domain-containing protein. J Cell Biol 145, 539–549.

Tan, B., Yatim, S., Peng, S., Gunaratne, J., Hunziker, W. and Ludwig, A. (2020). The Mammalian Crumbs Complex Defines a Distinct Polarity Domain Apical of Epithelial Tight Junctions. Curr Biol 30, 2791–2804 e2796.

Teo, G., Liu, G., Zhang, J., Nesvizhskii, A. I., Gingras, A. C. and Choi, H. (2014). SAINTexpress: improvements and additional features in Significance Analysis of INTeractome software. J Proteomics 100, 37–43.

Thomas, P. D., Ebert, D., Muruganujan, A., Mushayahama, T., Albou, L. P. and Mi, H. (2022). PANTHER: Making genome-scale phylogenetics accessible to all. Protein Sci 31, 8–22.

Tyanova, S., Temu, T. and Cox, J. (2016). The MaxQuant computational platform for mass spectrometry-based shotgun proteomics. Nat Protoc 11, 2301–2319.

UniProt, C. (2019). UniProt: a worldwide hub of protein knowledge. Nucleic Acids Res 47, D506–D515.

van den Goor, L. and Miller, A. L. (2022). Closing the gap: Tricellulin/alpha-catenin interaction maintains epithelial integrity at vertices. J Cell Biol 221.

Van Itallie, C. M., Aponte, A., Tietgens, A. J., Gucek, M., Fredriksson, K. and Anderson, J. M. (2013). The N and C termini of ZO-1 are surrounded by distinct proteins and functional protein networks. J Biol Chem 288, 13775–13788.

Van Itallie, C. M., Tietgens, A. J., Aponte, A., Fredriksson, K., Fanning, A. S., Gucek, M. and Anderson, J. M. (2014). Biotin ligase tagging identifies proteins proximal to E-cadherin, including lipoma preferred partner, a regulator of epithelial cell-cell and cell-substrate adhesion. J Cell Sci 127, 885–895.

Van Itallie, C. M., Tietgens, A. J., Krystofiak, E., Kachar, B. and Anderson, J. M. (2015). A complex of ZO-1 and the BAR-domain protein TOCA-1 regulates actin assembly at the tight junction. Mol Biol Cell 26, 2769–2787.

Wang, A., Dunn, A. R. and Weis, W. I. (2022). Mechanism of the cadherin-catenin F-actin catch bond interaction. eLife 11.

Wohlgemuth, S., Kiel, C., Kramer, A., Serrano, L., Wittinghofer, F. and Herrmann, C. (2005). Recognizing and defining true Ras binding domains I: biochemical analysis. J Mol Biol 348, 741–758.

Yamamoto, T., Harada, N., Kano, K., Taya, S., Canaani, E., Matsuura, Y., Mizoguchi, A., Ide, C. and Kaibuchi, K. (1997). The Ras target AF-6 interacts with ZO-1 and serves as a peripheral component of tight junctions in epithelial cells. J Cell Biol 139, 785–795.

Yang, Z., Zimmerman, S., Brakeman, P. R., Beaudoin, G. M., 3rd, Reichardt, L. F. and Marciano, D. K. (2013). De novo lumen formation and elongation in the developing nephron: a central role for afadin in apical polarity. Development 140, 1774–1784.

Yap, A. S., Duszyc, K. and Viasnoff, V. (2018). Mechanosensing and Mechanotransduction at Cell-Cell Junctions. Cold Spring Harb Perspect Biol 10, a028761.

Zaessinger, S., Zhou, Y., Bray, S. J., Tapon, N. and Djiane, A. (2015). Drosophila MAGI interacts with RASSF8 to regulate E-Cadherin-based adherens junctions in the developing eye. Development 142, 1102–1112.

Zhadanov, A. B., Provance, D. W., Jr., Speer, C. A., Coffin, J. D., Goss, D., Blixt, J. A., Reichert, C. M. and Mercer, J. A. (1999). Absence of the tight junctional protein AF-6 disrupts epithelial cell-cell junctions and cell polarity during mouse development. Curr Biol 9, 880–888.

Zhou, H., Xu, Y., Yang, Y., Huang, A., Wu, J. and Shi, Y. (2005). Solution structure of AF-6 PDZ domain and its interaction with the C-terminal peptides from Neurexin and Bcr. J Biol Chem 280, 13841–13847.

